# Shared landscapes, shared risk: *Echinococcus multilocularis* infection in coyotes is linked to agricultural land and altered green space in a major U.S. metropolitan area

**DOI:** 10.64898/2026.10.16.759553

**Authors:** Laurel E.K. Serieys, Leyi Wang, Shane McKenzie, Chris L. Anchor, Stanley D. Gehrt

## Abstract

Alveolar echinococcosis (AE), caused by the tapeworm *Echinococcus multilocularis*, is among the most lethal zoonotic helminthic diseases affecting humans. Historically rare in North America, human and canine AE cases have risen sharply over the past decade, coinciding with the spread of a more virulent European-type haplotype and the emergence of coyotes (*Canis latrans*) as a principal definitive host in and around urban centers. Despite growing recognition of *E. multilocularis* as an emerging public health concern, the host- and landscape-level factors that concentrate infection risk within urban environments remain poorly understood. We used monitoring data from the Chicago metropolitan area to assess *E. multilocularis* prevalence and test demographic and land cover predictors of infection among urban coyotes. Fecal samples from 85 coyotes were screened via real-time PCR, and land cover association with *E. multilocularis* infection within individual home ranges or capture-based buffers was investigated using generalized linear models. Overall, 27.1% of coyotes were infected. Infection status was unrelated to age or sex, but coyotes with a greater proportion of agricultural land or altered open land cover, such as golf courses, cemeteries, and manicured parks, within their home ranges had significantly higher odds of infection (OR_agriculture_ = 2.84; OR_altered open spaces_ = 2.07). These landscapes are also heavily used by people and domestic pets, suggesting elevated potential for zoonotic spillover in human-dominated areas rather than natural or forested habitat. Our findings identify specific, human-accessible landscapes that sustain *E. multilocularis* transmission in a major U.S. metropolitan area and provide a framework for prioritizing surveillance, targeted anthelmintic intervention, and public health outreach in expanding urban coyote populations across North America.

## Introduction

*Echinococcus multilocularis* is a small tapeworm of canids that can cause alveolar echinococcosis (AE), one of the most lethal zoonotic helminthic diseases affecting humans [1]. AE develops when the larval stage of the parasite proliferates within host tissue, most often the liver, in a slow-growing, tumor-like pattern that is fatal in a large proportion of untreated cases [2]. *E. multilocularis*, found in tropical, temperate, and arctic biomes [3], is broadly distributed across the Northern Hemisphere, occurring throughout large areas of Europe, Asia, and North America [4,5]. The geographic range and prevalence of this cestode appears to be increasing in part due to anthropogenic influences such as increased globalization of animals and altered human-animal interfaces [5]. AE is considered to be a significant emerging zoonotic [1,5,6]. AE is thus of growing global concern due to the geographic spread of the parasite, its rising prevalence in animals within endemic areas, and the absence of control measures to prevent infection [1].

The parasite is maintained through an indirect, predator-prey transmission cycle involving a definitive host and an intermediate host. Adult tapeworms reside in the small intestine of wild canid definitive hosts, principally red foxes (*Vulpes vulpes*) and arctic foxes (*Vulpes lagopus*) in Europe and Asia, and coyotes (*Canis latrans*), foxes (*Vulpes spp.*), and wolves (*Canis lupus*) in North America [4]. *Echinococcus* tapeworms release eggs that are passed in host feces and that can remain infective in the environment for extended periods, especially under cool, moist conditions [2,4,7]. Eggs are ingested by small mammal intermediate hosts, particularly rodents such as voles [8]. Once ingested, the egg hatches in the intermediate host’s gut, and the released larva penetrates the intestinal wall and migrates to the liver, where it develops into the infective stage [4,9]. The cycle is completed when an infected intermediate host is consumed by a definitive host (e.g. a wild canid), allowing the tapeworms to develop within the canid intestine [10]. Humans are dead-end intermediate hosts, becoming infected incidentally after ingesting eggs through contaminated food or water, direct contact with infected animals, or environmental exposure [1].

Although *E. multilocularis* has long been recognized as a public health concern in Europe and Asia, locally-acquired human infection in North America was, until recently, exceedingly rare. Only a small number of confirmed cases were documented outside a single endemic focus on St. Lawrence Island, Alaska, prior to the early 2010s [11]. This pattern has shifted markedly over the past decade: since 2013, at least 17 locally acquired human AE cases have been confirmed in Alberta, Canada [12]. Additional cases in people and infections in domestic dogs have also been reported in Ontario [13] and, more recently, in the northeastern United States [14]. This rise in reported human and canine cases has coincided with the detection of a novel, European-type strain of *E. multilocularis* in wildlife across parts of western Canada. The European-type haplotype appears more virulent and more likely to progress to clinical AE than the previously circulating North American strain [15]. The European-type strain is now the most common endemic haplotype in North America [16,17]. This apparent expansion of both the parasite’s geographic range and pathogenic potential underscores the need for a better understanding of the wildlife reservoirs sustaining *E. multilocularis* transmission in North America, and the risk factors associated with infection.

In North America, *E. multilocularis* was historically confined to sylvatic transmission cycles in the north-central United States and Canada. An early survey in the 1960s reported prevalence as low as 4.1% in coyotes in North Dakota [18]. Since then, however, coyotes have emerged as a principal definitive host [19], maintaining the parasite in and around urban centers across the continent [11]. Infection prevalence in urban coyote populations may be high. In Edmonton, Alberta, Canada, prevalence was 65% [20,21]. Over the past few decades, there has been little focus on *Echinococcus spp.* tapeworm research in the contiguous United States [22]. Therefore, few figures are available for infection prevalence in wild canids across the United States. In the central United States, however, prevalences in coyote studies were found to be 19–44% [23,24]. Most recently, detections have extended the parasite’s documented range into the Pacific Northwest [25] and New York [26]. This expansion is of particular concern because coyotes have become widespread and abundant across North American cities, where they persist in close proximity to humans and domestic pets [27]. Urban and suburban coyotes rely heavily on small rodents, especially voles (important intermediate hosts for *E. multilocularis*), as dietary staples [27,28]. Coyotes are therefore well suited to sustain the sylvatic transmission cycle even within highly developed landscapes. Yet despite these signs of an expanding and ecologically well-supported transmission cycle, the specific factors that determine where infection risk is concentrated within urban landscapes remain poorly understood.

As *E. multilocularis* gains recognition as an emerging public health concern in urban North America, identifying the hosts and landscapes that sustain transmission is essential for anticipating where human and canine exposure risk is greatest. Here, we used monitoring data to assess *E. multilocularis* prevalence and generalized linear models to test host-level (age, sex) and landscape-level predictors of *E. multilocularis* infection among urban coyotes sampled across the Chicago metropolitan area (CMA). The CMA is one of the largest and most heterogeneous urban regions in North America and encompasses a mosaic of development, agriculture, forest preserves, and managed open space. The CMA offers an opportunity to evaluate how land use within individual coyote home ranges relates to infection status. By identifying which host traits and landscape features are most strongly associated with infection, this study provides a framework for anticipating *E. multilocularis* risk in other expanding urban coyote populations and informs the surveillance and management strategies needed to curb this growing zoonotic threat.

## Methods

### Study area

We conducted this study as part of the Urban Coyote Research Project (Urbancoyoteresearch.com), an ongoing long-term research initiative investigating coyote ecology within the Chicago Metropolitan Area (CMA) since 2000. The CMA represents one of the largest metropolitan regions in North America spanning 28,120 km^2^ with a human population exceeding 9 million. Coyotes are distributed throughout the CMA. Land uses include low-, medium- and high-density development, commercial areas, forest preserves, grassland, altered open areas (e.g. cemeteries, golf courses, and manicured parks), agricultural areas (pasture and crops), shrubby areas and wetlands (Figure 1). Winter temperatures and precipitation averaged −0.8°C and 15.6 cm, while summer temperatures and precipitation averaged 23.6°C and 31.3 cm (National Weather Service, Chicago, IL Forecast Office). Besides coyotes, other native fauna of importance in the CMA include both definitive and potential intermediate *E. multilocularis* hosts (Deplazes et al., 2017; Romig and Wassermann, 2024) such as red fox (*Vulpes vulpes*), voles (*Microtus spp.*), white-footed mice (*Peromyscus* spp.), muskrat (*Ondatra zibethica*), and eastern cottontail (*Sylvilagus floridanus*). Non-native fauna include domestic cats (*Felis catus*) and domestic dogs (*Canis familiaris*) [28].

**Figure 1.**
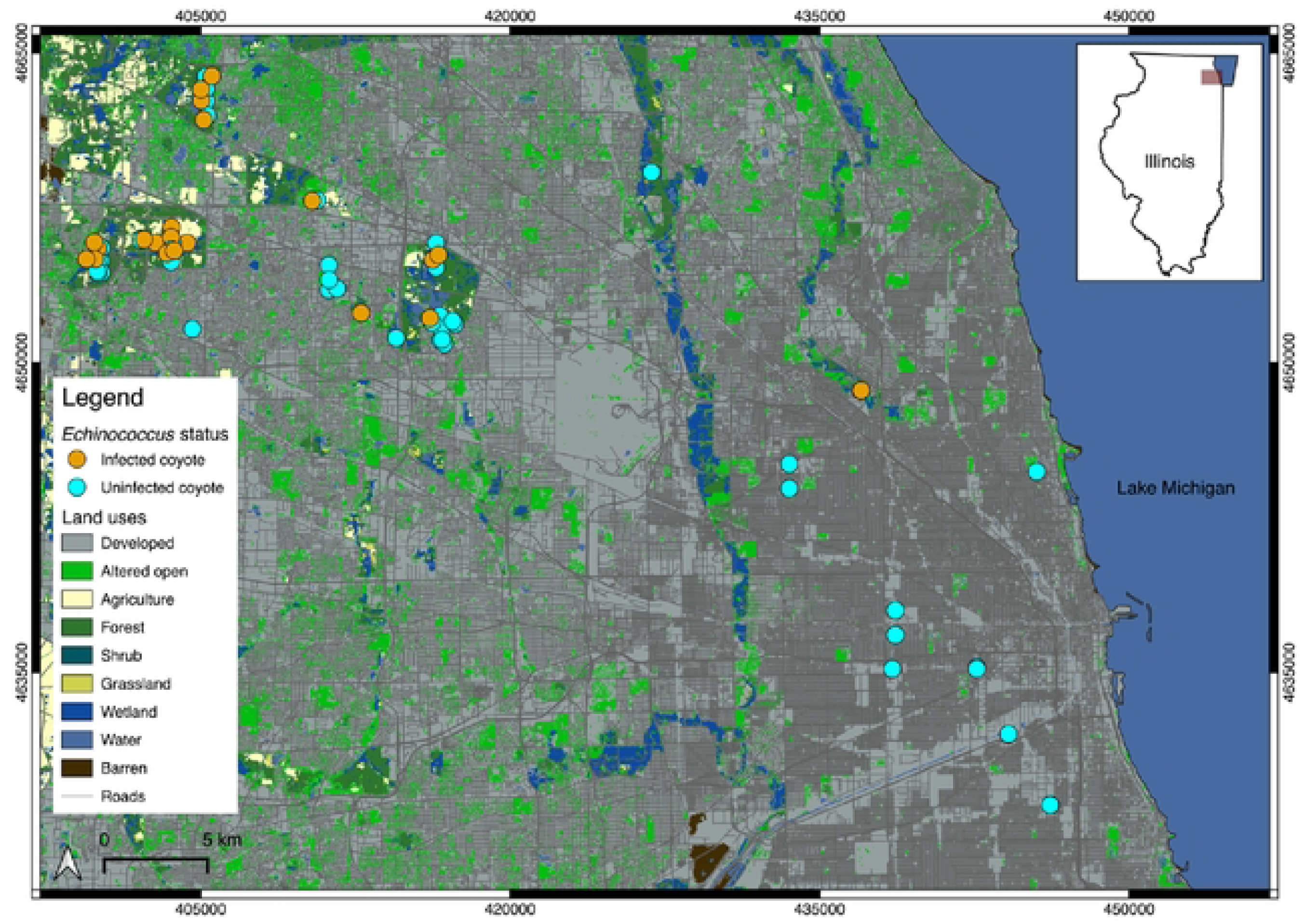
Map of the study area and the locations of individuals infected with *Echinococcus multilocularis*.

### Capture and sample collection

Coyotes were captured between 2016–2021 in the central and northwestern portion of the CMA. Captures primarily occurred in forest preserves, golf courses, small woodland parks, or in abandoned lots. Although captures were conducted throughout the year, they predominantly occurred in winter and early spring. Coyotes were captured using neck cable restraint devices and foothold traps [29]. Once captured, individuals were transported in a carrier to a research laboratory. Coyotes were sedated with 2.5 mg/kg Telazol (Fort Dodge Animal Health, Fort Dodge, IA, USA). Following immobilization, each coyote was sexed and aged based on reproductive condition and tooth wear [30]. Age classes included pup (<1 year), subadult (1–2 years), and adult (>2 years). Fecal samples were collected at the time of capture and stored at −80°C until testing. All coyotes were ear-tagged and fitted with very-high-frequency radio (VHF) collars (Advanced Telemetry Systems, Isanti, MN, USA) or with GPS-collars (Lotek Wireless, Newmarket, Ontario, Canada). After recovery from sedation, individuals were released at the capture locations.

### Ethics statement

All capture and handling procedures followed the guidelines of the American Society of Mammalogists [31]. Coyote captures were approved by the Ohio State University (IACUC ID: 2013A00000012-89 R1) and by the Illinois Department of Natural Resources (permit no.: IDNR W17.0122).

### Echinococcus testing

Fecal samples were submitted to the Veterinary Diagnostic Laboratory at the University of Illinois Urbana Champaign for *E. multilocularis* screening. Feces were swabbed with a cotton-tipped wooden applicator, and the swab tip was placed in a 1.5 mL microcentrifuge tube containing 1,000 µL of phosphate-buffered saline (PBS), agitated to release adherent material, and then removed. The suspension was vortexed for 20 s and centrifuged at 6,000 × *g* for 2 min at room temperature to pellet debris. A 200 µL aliquot of the supernatant was used for nucleic acid extraction with the MagMAX Pathogen RNA/DNA Kit (Thermo Fisher Scientific, Waltham, MA, USA) on a KingFisher Flex automated purification system (Thermo Fisher Scientific) according to the manufacturer’s instructions. Extracted nucleic acids were stored at −80 °C until analysis.

For real-time PCR detection of *E. multilocularis*, primers and a probe targeting the mitochondrial *rrnL* gene were adapted from a published protocol [32]. The primer and probe sequences were as follows: Em-rrn forward primer, 5′-CTGTGATCTTGGTGTAGTAGTTGAGATTT-3′; Em-rrn reverse primer, 5′-GGCTTACGCCGGTCTTAACTC-3′; and Em-probe, 5′-Cy5-TGGTCTGTT-BHQ-CGACCTTTTTAGCCTCCAT-IAbRQ-3′. Each 25 µL PCR reaction contained 6.25 µL of TaqPath 1-Step RT-qPCR Master Mix, 1 µL of 10 µM forward primer, 1 µL of 10 µM reverse primer, 1 µL of 5 µM probe, 5 µL of extracted nucleic acid template, and 10.75 µL of nuclease-free water. Real-time PCR was performed on a Bio-Rad CFX Opus 96 Real-Time PCR System (Bio-Rad, Hercules, CA, USA) under the following conditions: UNG incubation at 25 °C for 2 min, reverse transcription at 48 °C for 10 min, initial denaturation at 95 °C for 10 min, and 40 cycles of 95 °C for 15 s and 60 °C for 45 s. Results were analyzed using CFX Maestro software (Bio-Rad, Hercules, CA, USA, version 1.0).

### Home range and buffer estimations

Relocations of all VHF-collared coyotes were recorded, on average, 8–14 times per month with 3–4 day locations and 5–10 night locations. GPS-collars recorded locations ranging from 0.25–7.25 hours. For VHF-collared animals, we estimated home ranges only if they had a minimum of 50 GPS locations. For GPS-collared individuals, we used all available GPS locations for each coyote to estimate home ranges. Home ranges were quantified as a 95% autocorrelated kernel density estimates (95% aKDE) using the ‘amt’ package [33] in R [34]. This approach applies a continuous-time movement modelling framework [35] and corrects for autocorrelation among location fixes and yields unbiased home range estimates despite irregular sampling intervals [36]. For each individual, candidate continuous-time movement models (isotropic and anisotropic IID, OU, and OUF) were fit to the location data and compared via AIC to identify the best-supported model [36]. The best-supported model was then used to compute the 95% AKDE. We excluded individuals with an effective degrees of freedom for area (DOF_area_) below 15, since a low DOF_area_ reflects inadequate data relative to the position autocorrelation timescale and yields unreliable home range estimates [37]. In cases where GPS- or VHF-data were unavailable for individuals, we drew circular buffers around the capture location of individuals sized according to the median 95% aKDE home range estimates for each demographic group: adult females 10.3 km^2^ (n = 90), subadult females 16.6 km^2^ (n = 69), pup females 11.5 km^2^ (n = 29), adult males 11.2 km^2^ (n = 141), subadult males 11.1 km^2^ (n = 84), and pup males 8.1 km^2^ (n = 27).

### Landscape covariates

For land cover predictors, we obtained annual land cover classifications within home ranges or buffers for 2016–2021 from the Annual National Land Cover Database (NLCD) product (U.S. Geological Survey, 2024), pulling data from the annual NLCD GIS layer to match the year of coyote capture. From these layers, we calculated the proportion of development, altered open areas, agriculture, water, wetlands, shrubs, forest, and grassland. We also calculated the average normalized difference vegetation index (NDVI; European Space Agency, 2024, Sentinel-2; 10 m resolution) within the home ranges or buffers, with each NDVI layer year matching the year of sample collection. Because *E. multilocularis* thrives in moist environments, we extracted information about soil and vegetation moisture. We first characterized soil water-holding capacity using SoilGrids 2.0 [38], a machine learning–based global soil property product available at 250 m resolution. We obtained modeled volumetric water content (cm³/cm³) at three suction levels across six standard depth intervals, and calculated a thickness-weighted average of each suction level across the 0–30 cm depth range (topsoil/rooting zone). We then derived plant available water capacity (PAWC) as the difference between water content at field capacity and at the wilting point, representing the fraction of soil water accessible to plant roots, and used PAWC as our soil moisture–related covariate. Finally, as an additional moisture proxy, we extracted the average normalized difference moisture index (NDMI; European Space Agency, 2024, Sentinel-2; 10 m resolution) which measures vegetation moisture content and can help identify wetter or more saturated areas within the landscape.

We found that average NDVI and the proportion of development, forest, grassland and wetlands were all colinear. Average NDMI was also colinear with the proportion of development. Given that one of our primary interests was whether development was a spatial predictor of infection, we dropped average NDVI, average NDMI, and the proportion of forest and wetlands from our analyses. All other landscape covariates tested in models (proportion of development, altered open, agriculture, and shrubs, and PAWC) had correlation values of r ≤ |0.64|. Variance inflation factors (VIF) for covariates in the best-fit model were <2 [39]. All landscape covariates were scaled with a mean = 0 and standard deviation = 1.

### Statistical analyses

We used generalized linear models (GLMs) with a binomial error distribution and logit link to assess associations between *E. multilocularis* infection status (infected/uninfected) and land cover and host-level covariates. Candidate predictors included six scaled land cover variables and two host covariates: age class and sex.

Model selection was performed using an information-theoretic approach. Starting from a global model with all land cover and host covariates, all possible subsets of covariates were evaluated using the dredge function in the R package ‘MuMIn’ [40]. Models were ranked by the corrected Akaike Information Criterion (AICc), which adjusts for small sample size relative to the number of parameters [41,42]. Multiple models within 2 AICc units of the top-ranked model (ΔAICc<2) were produced and the most parsimonious one was selected. Coefficients from this model are reported as odds ratios with 95% confidence intervals.

Goodness of fit for the best-supported model was assessed using simulation-based scaled residuals implemented in the R package ‘DHARMa’ [43]. Residuals were simulated from the best-fit model (n = 1000 simulations) and tested for deviations from uniformity, over/underdispersion, and excess outliers relative to the expected distribution under the model.

All analyses were performed in R statistical software, version 4.3.2 [34].

## Results

### Sampling

We collected fecal samples from 85 individuals. We sampled 15 adult females, 23 adult males, 8 subadult females, 8 subadult males, 15 pup females, and 16 pup males. Among the 85 samples, 27.1% of individuals (23/85) were infected with *E. multilocularis*. For 37 individuals, we used 95% aKDE home range estimates, and for 48 individuals we used buffers to extract landscape covariates.

### Spatial predictors of infection

We detected only two landcover predictors of infection (Figure 2), altered open areas (Estimate = 0.73, 95% CI: 0.12–1.34) and agricultural areas (Estimate = 1.04, 95% CI: 0.43–1.65). Specifically, coyotes with a greater proportion of altered open land cover (e.g., cemeteries, golf courses, manicured parks) within their home range had higher odds of *E. multilocularis* infection (OR = 2.07, 95% CI: 1.17–4.06), as did coyotes with a greater proportion of agricultural land (OR = 2.84, 95% CI: 1.60–5.52). As the proportion of altered open area or agricultural land cover increased within home ranges or buffers, the probability of infection with *E. multilocularis* also increased (Figure 2). The mean proportion of altered open areas in home ranges or buffers was 9.1 % (SD = 7.1 %). The mean proportion of agricultural areas was 9.5% (SD = 10.8 %). We found no demographic predictors of infection nor associations with moisture indices.

**Figure 2.**
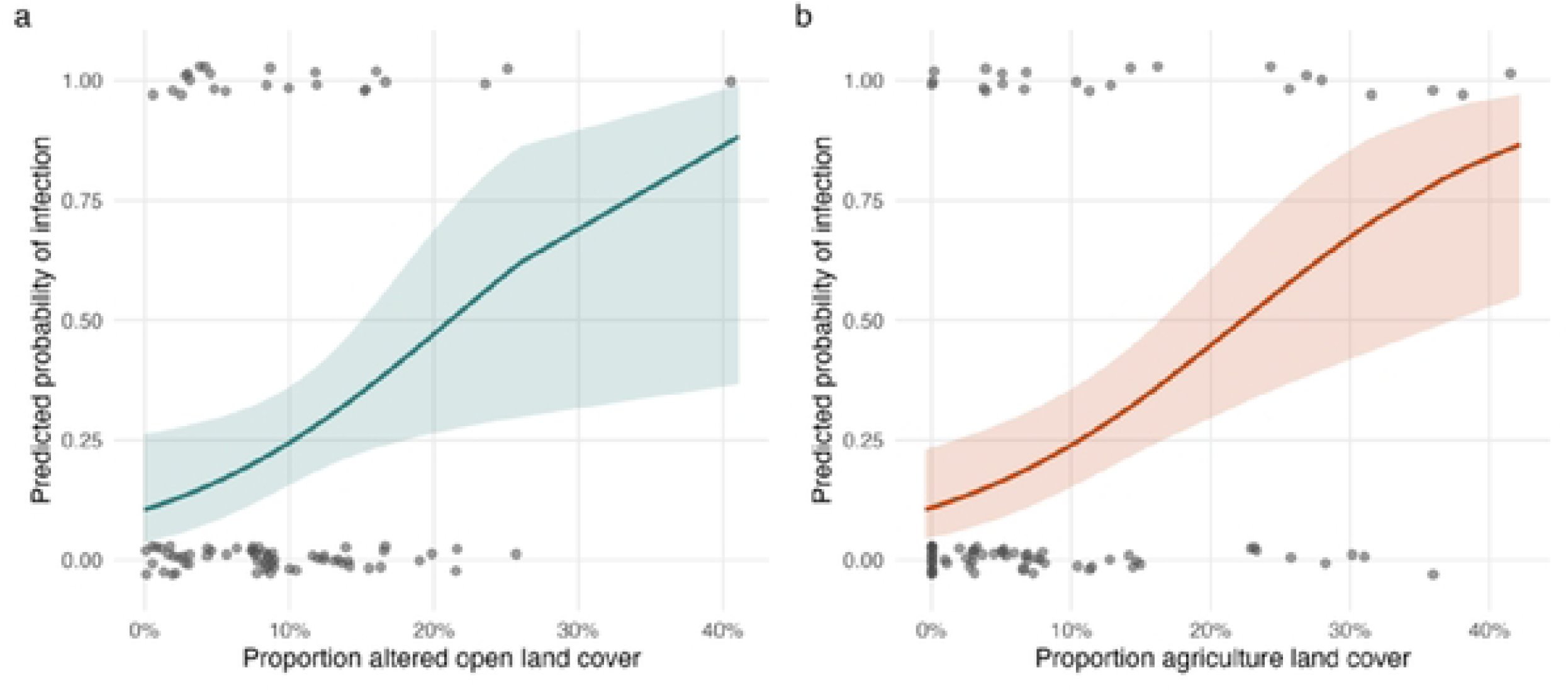
Plots displaying the predicted probability of infection as the proportion of altered open land cover (a) and proportion of agricultural land cover (b) in home ranges or buffer zones increase.

### Model goodness of fit

All tests indicated no evidence of lack of fit: residuals did not deviate from uniformity (Kolmogorov-Smirnov test, D = 0.06, p = 0.93), there was no evidence of over- or underdispersion (dispersion = 1.02, p = 0.89), and no outliers were detected (0 of 85 observations, p = 1).

## Discussion

We investigated *E. multilocularis* infection prevalence and demographic and land cover predictors of infection in coyotes of the CMA. We found that 27.1% of urban coyotes sampled across the CMA were infected with *E. multilocularis*, a prevalence broadly consistent with the 19–44% previously reported in the United States [23–25]. These results are, however, markedly lower than the 65% prevalence reported in urban coyote populations in Edmonton, Alberta, Canada [20,21]. Infection status was unrelated to coyote age class or sex. Instead, we identified two spatial predictors of infection: coyotes with a greater proportion of agricultural land and a greater proportion of altered open area within their home ranges had significantly higher odds of infection. The proportion of agricultural land in home ranges or buffers showed the strongest effect. Nevertheless, because these landscape predictors are human-use areas, even this comparatively moderate prevalence of *E. multilocularis* infection is enough to sustain a risk of spillover into domestic dogs and humans [13].

Explicit studies examining land cover predictors of *Echinococcus* infection are relatively few, although past literature has similarly linked agricultural and grassland-dominated landscapes to elevated *E. multilocularis* transmission in wildlife. In eastern France and central China, AE prevalence and infection in small mammal intermediate hosts were strongly associated with the proportion of grassland cover [44]. Similarly, in a survey of definitive hosts in the Baltic states, there was a significant positive association between agricultural land cover and *E. multilocularis* prevalence in raccoon dogs (*Nyctereutes procyonoides*). A study of coyotes in the southern Ontario region of Canada found a negative association between the proportion of natural area in buffer zones and infection, but found no relationship with the proportion of agricultural area [45]. In Germany, infected red foxes contained a greater proportion of pasture and a decreased proportion of forest in their buffer zones [7]. Our finding that altered open areas, a land-cover class encompassing golf courses, cemeteries, and manicured parks, were also positively associated with infection is notable. These areas are typically dominated by mown or maintained turf and grassy vegetation within, or adjacent to, residential and commercial development. Functionally, with respect to *E. multilocularis* infection risk, however, they may act like grassland-dominated systems that have been linked with *E. multilocularis* infection in host species and in humans in both China and France [44]. Collectively, these studies combined suggest that anthropogenically altered landscapes tend to elevate *E. multilocularis* transmission risk relative to natural or forested land although the specific land-cover types implicated vary by region and host species.

The mechanism underlying the association with these landcovers and *E. multilocularis* infection is likely rooted in prey availability within the different land covers. Predator-prey dynamics play a central role in shaping the transmission intensity of trophically transmitted parasites [46]. Deer mice (*Peromyscus maniculatus*), meadow voles (*Microtus pennsylvanicus*), house mice (*Mus musculus*), and the muskrat are documented intermediate hosts in North America [8,18,22] although others likely exist, especially among arvicoline rodents [8]. Deer mice, meadow voles, house mice and muskrats all occur in the CMA [47] and meadow voles and white-footed mice constituted 88% of all small rodents trapped during a coyote diet study in the CMA [28]. Deer mice and meadow voles are associated with open grassy habitats, cropland edges [48], and lawns and golf courses [49], the type of land uses associated with *E. multilocularis* infection in this study. Coyotes are opportunistic but rodent-focused predators [27], particularly of voles in some areas including Chicago [28,50]. Agricultural and altered open landscapes that may support abundant vole populations likely increase both the coyote’s dietary encounter rate with infected intermediate hosts and, in turn, its probability of acquiring infection [46]. This relationship was demonstrated in urban coyotes in Calgary. Transmission of the parasite to coyotes was mediated by prey selection, with a preference for meadow voles elevating coyotes’ encounter rate with the parasite beyond what would be expected from the rodent community’s composition alone [50]. Notably, the same research group documented that *E. multilocularis* prevalence varied widely (5–61%) among Calgary’s urban parks, suggesting that fine-scale differences in habitat structure and prey community composition, and not coarse land-cover classification alone, shape local transmission intensity [51]. If a comparable relationship holds in Chicago, land cover categories associated with elevated rodent, and particularly vole, abundance may better predict local *E. multilocularis* infection risk than land use alone.

Contrary to our initial expectation, soil water-holding capacity (PAWC) did not emerge as a predictor of infection, despite the ecological rationale for including a measure of soil moisture in our models. Further, infected red foxes in Germany were associated with areas of high soil humidity [7]. *E. multilocularis* eggs are known to be highly sensitive to desiccation and persist longest under cool, moist conditions [2,4,7,52]. There may be several reasons for our null result. First, the spatial and temporal resolution of our moisture covariate may not adequately capture the microhabitat conditions that actually promote egg survival. Because egg viability is extremely sensitive to elevated temperatures and to desiccation, it is likely determined by microhabitat-scale conditions (e.g., shade, humidity, soil moisture) [52]. On the other hand, we used PAWC to characterize average moisture conditions across an entire home range or buffer. Second, moisture conditions favorable to egg survival may be a necessary but not limiting factor across much of the CMA; if ambient moisture rarely falls low enough to meaningfully reduce egg viability anywhere within our study area, especially given the region’s temperate, humid-continental climate (National Weather Service, Chicago, IL Forecast Office), then variation in moisture would have little power to explain variation in coyote infection. Third, infection risk for a definitive host such as the coyote is influenced primarily by the probability of ingesting an infected intermediate host [50], not directly by the egg’s environmental survival. Land-cover variables that index prey habitat and rodent abundance may therefore capture the dominant transmission pathway more effectively than variables that characterize abiotic egg persistence alone. Under this interpretation, the significance of agricultural and altered open land cover, and the null result for moisture, together point toward prey availability rather than environmental egg survival as the more proximate driver of spatial variation in coyote infection risk within this landscape.

Previous studies have found different results regarding the association of *E. multilocularis* infection and demographic factors, but we did not observe any demographic risk factors in our study. Sex was not found to be a predictor of infection in coyotes in Edmonton, Alberta, Canada [20] or across the southern Ontario region [45] which may be the result of similarly sized home ranges among the coyote demographic groups (e.g., this study). If, like some other carnivore species, males had substantially larger home ranges and further ranging movements (e.g. bobcats (*Lynx rufus*; [53] or mountain lions (*Puma concolor*; [54]), they may be more at risk to encounter an infected intermediate host. Age class was also not a predictor of infection. In red foxes and raccoon dogs of Latvia, there was no association between age and infection prevalence or intensity of infection [55]. However, pup (<1 year) coyotes from Calgary, Alberta had a higher prevalence of *E. multilocularis* infection compared to adults [56]. Juvenile red foxes were also significantly more heavily infected with *E. multilocularis* tapeworms compared with adults in Geneva, Switzerland [57] and in Zurich, Switzerland [58]. These findings are surprising considering that as animals age, they have an increased risk of exposure to the *Echinococcus* tapeworm [59]. Our samples were well stratified across differing age groups but as a result, we had relatively few samples per age group. Thus, we recommend that future studies use a larger sample size across multiple demographic groups to better test whether age is indeed a demographic risk factor in coyotes.

The landscape associations we found carry direct implications for human and domestic dog exposure risk. In a case-control study of AE patients in Germany, farming and residential proximity to agricultural fields were among the strongest risk factors for human infection [60]. Specifically, 65% of human AE cases were associated with farming. A similar link between farming and human AE cases was also found in Central Europe [61]. This association presumably reflects a greater likelihood of environmental contamination with infective eggs in open, cultivated landscapes [60]. Our results, together with this broader literature, suggest that the same agricultural and grassy altered open areas that elevate coyote infection risk are also plausible sites of human environmental exposure [62]. This is particularly relevant in a landscape such as the CMA, where agricultural land and manicured green spaces are frequently interspersed with, or immediately adjacent to, residential neighborhoods. These spaces may also be heavily used by people and by off-leash or free-roaming pets. Domestic dogs, which are definitive hosts typically infected by consuming intermediate hosts [2], are thought to contribute significantly to the *E. multilocularis* life cycle in urban areas of Europe [62] and some pastoral areas of central Asia [63,64]. One *E. multilocularis* transmission route to humans is simply handling a dog with *Echinococcus* eggs on its fur or snout [60]. In China and Alaska, high prevalences of *E. multilocularis* were found in domestic dogs and were thought to be the primary sources of infections in humans in villages [65,66]. Therefore, the use of agricultural and especially altered open areas by domestic dogs in Chicago may feed the *E. multilocularis* life cycle and also put their human owners at risk.

*Echinococcus* is a silvatic parasite involving multiple wild animal hosts, and therefore complete elimination of the tapeworm from the landscape is impractical [2]. However, monthly anthelmintic baiting of red foxes with praziquantel in Zurich, Switzerland was found to not only reduce *E. multilocularis* infection prevalence in the foxes, but it also reduced the prevalence in the local intermediate host, the European water vole (*Arvicola terrestris*) [67]. Similarly, a study in an Alaskan village where domestic dogs were routinely treated with praziquantel led to a lower prevalence of *E. multilocularis* in locally trapped northern voles (*Microtus oeconomus*) [68]. Multiple studies have shown that the distribution of *E. multilocularis* biomass can be highly aggregated whereby a few highly infected foxes are responsible for most of the environmental egg contamination [57,62,69]. For example, 8% of 388 red foxes carried 72% of the total biomass of *E. multilocularis* in Zurich, Switzerland [69]. This aggregation suggests that control efforts targeting high-risk habitats with anthelmintic baits could have an outsized effect on reducing the prevalence in the environment if highly infected individuals consume the bait. In the CMA, if an anthelmintic campaign were implemented, then prioritizing praziquantel baiting in high-risk habitats could meaningfully reduce the amount of egg-shedding in the environment. Coyotes captured in the CMA could also be treated with praziquantel at the time of capture. However, in the studies where baiting or regular anthelmintic treatment occurred, the tapeworm was still not entirely eliminated from the environment and so these measures may reduce the occurrence of the tapeworm in the CMA but not completely eliminate it.

Together, these findings offer a practical starting point for targeting *E. multilocularis* surveillance and risk communication in urban landscapes like the CMA. Because infection risk was concentrated in agricultural and altered open land covers rather than distributed uniformly across the urban matrix, monitoring and public health messaging can be prioritized in these areas rather than applied evenly across a metropolitan region as large and heterogeneous as Chicago. Given that these same land covers are heavily used by people and domestic dogs, targeted interventions, such as periodic PCR fecal screening of domestic dogs [17], routine anthelmintic treatment of domestic dogs [68], signage discouraging off-leash access in golf courses, cemeteries, and agricultural areas, and outreach to farmers and land managers about hand hygiene and produce washing, may complement broader regional surveillance. As *E. multilocularis* continues to expand its range and pathogenicity in North America [5,15], identifying the specific landscapes that sustain transmission will be essential for allocating limited surveillance resources where they are most likely to reduce zoonotic spillover risk.

## Acknowledgements

Financial, logistical support, and facilities were provided by Cook County Animal and Rabies Control, Max McGraw Wildlife Foundation, and the Forest Preserves of Cook County. We thank the many students and field technicians that assisted with fieldwork.

